# Loss of starvation‑and‑light sporulation trigger in Myxomycetes *Physarum roseum*

**DOI:** 10.1101/2025.05.10.652879

**Authors:** Mana Masui, Phillip K. Yamamoto, Nobuaki Kono

**Author notes:** Author for correspondence: Nobuaki Kono.

## Abstract

Myxomycetes are unicellular amoebozoans that form fruiting bodies to reproduce (sporulation). In the model species *Physarum polycephalum*, this morphogenesis is triggered when starvation is followed by starvation-plus-light cue has been considered broadly conserved throughout *Physarum*. Recent observations of congeners that fail to sporulate under the same conditions have raised doubts about this assumption and prompted tentative taxonomic reconsideration. Because comparable starvation and light tests are scarce for other *Physarum* species, their phenotypes and molecular mechanisms remains unclear. Consequently, we investigated *Physarum rigidum* and *Physarum roseum* under starvation and light. Four of six *P. rigidum* plasmodia sporulated by day 6, whereas *P. roseum* did not sporulate within seven days. RNA-seq of *P. roseum* across nutrient-rich/starved and dark/light conditions showed that differential expression was driven chiefly by nutrition; light caused only minor changes and did not elicit the transcriptional program characteristic of *P. polycephalum* sporulation. The photoreceptor genes that drive sporulation in *P. polycephalum* were not detected in *P. roseum*, and 92 candidate photoreceptor genes showed no significant regulation. These findings indicate that *P. roseum* responds only minimally to light stimulation, and that the starvation-plus-light trigger is not universally retained within *Physarum*.

## Introduction

Myxomycetes (Slime mould) are multinucleate unicellular organisms belonging to the Amoebozoa and known to display diverse morphologies throughout their life cycle [1]. The plasmodium moves at approximately 1 cm / h [2] and typically inhabits moist, dark microhabitats such as decaying wood or leaf-litter, where it preys on bacteria and fungi [3]. When exposed to environmental stressors such as starvation or light, the plasmodium either enters a metabolically inactive dormant state (sclerotium) or proceeds to form fruiting bodies. In the model Myxomycetes *Physarum polycephalum*, and *Didymium iridis*, starvation and light have been demonstrated to be indispensable cues for sporulation [4,5]. After more than three days of starvation preconditioning, *P. polycephalum* synthesises several photoreceptors, including phytochrome-like proteins that respond to wavelengths from the visible to the near-infrared range. A brief far-red pulse (∼700 nm) or several hours of continuous visible light then acts as a trigger, inducing sporulation [6.7]. Although plasmodia generally exhibit negative phototaxis [8], starvation can occasionally induce positive phototactic behaviour [9]. In the absence of light stimulation after starvation, the plasmodium bypasses sporulation and instead forms a sclerotium [10,11].

Fruiting-body induction by combined starvation and light stimuli is thought to be widely conserved, including across species of the genus *Physarum* [12,13]. Members of the genus also share core ecological traits, including similar microhabitats, large phaneroplasmodium, and prolific fruiting [14], and are widely employed as model organisms in Myxomycetes research. Recent molecular phylogenies, however, propose transferring several species formerly assigned to *Physarum* to other genera, indicating that the current taxonomy warrants re-evaluation [15].

Phenotypic data likewise reveal diversity: *Physarum nivale* grows only within an exceptionally narrow, low-temperature range [16], whereas our observations show that *Physarum roseum* thrives over a markedly broader temperature condition. In the inter-plasmodial allorecognition (Fusion and Avoidance) universally observed among Myxomycetes, heterospecific fusion is typically absent. In heterospecific encounters within the genus *Physarum*, not only is avoidance behaviour lacking, but the two plasmodia appear unable to recognize each other [17]. Critically, the sporulation trigger may differ among *Physarum*: *P. roseum* almost never forms fruiting bodies under conditions that reliably induce sporulation in *P. polycephalum* (starvation ≥ 3 days under dim natural light) [17].

Taken together, these findings indicate that ecological, physiological, and cultivation traits are likely to vary considerably even among closely related *Physarum* species. Because sporulation is the life-cycle stage under the strongest selective pressure, dissecting its interspecific variation is fundamental to understanding evolution within the genus. Nonetheless, detailed ecological, functional and molecular investigations of *Physarum* species other than *P. polycephalum* remain scarce, and their transcriptional responses to starvation and light are virtually unexplored.

To clarify the diversity of responses to the starvation and light cues that Induce sporulation, we will compare both phenotypic and intracellular reactions in plasmodia of *Physarum* species closely related to *P. polycephalum*. Phenotypically, we will examine behavioural responses to starvation and light stimulation in these congeners. Intracellular dynamics will be assessed by differential expression gene analysis: plasmodia will be cultured under factorial combinations of light versus dark and nutrient-rich versus starving conditions, and transcriptomic changes across treatments will be profiled by RNA-seq.

## Method

### (a) Samples

Plasmodia of *Physarum roseum* strain Ro1 were collected in 2018 during fieldwork at the Midori-no-Mori Museum, Saitama Prefecture, Japan, and plasmodia of *Physarum rigidum* strain Ri1 were obtained from the same locality in 2020. Both strains had already been taxonomically verified in a previous study [17] by DNA barcoding of the SSU rRNA region with two primer sets, together with morphological examination of sporocarp characters (peridium, spores, and capillitium), and were used herein as pre-identified reference strains.

### (b) Culture conditions

Plasmodia were maintained in 9 cm plastic Petri dishes containing 2% plain (nutrient-free) agar. Dishes were kept in an incubator at 23 °C and, under normal maintenance, were completely shaded from light. Each day at 16:00 the agar surface was moistened with sterile water, and rolled oats were supplied as a food source.

For RNA-seq, plasmodia of *P. roseum* and *P. rigidum* were cultured for three days under four factorial conditions combining light (dark vs. light) and nutrition (nutrient-rich vs. starved).

- Light treatment: dishes were exposed to diffuse daylight transmitted through a translucent panel together with ambient LED room lighting.
- Starvation treatment: no oats were provided; only sterile water was added at the daily watering time.

All other parameters (dish size, agar composition, incubation temperature, watering schedule) were identical to the maintenance conditions described above.

### (c) Light- and Starvation-Response Assay in *P. roseum* and *P. rigidum*

Plasmodia of *P. roseum* and *P. rigidum* were transferred to fresh plates of plain (nutrient-free) agar and cultured for up to seven days under conditions reported to induce sporulation in P. polycephalum [7, 18]. Cultures were exposed to diffuse daylight passing through a translucent panel together with ambient LED room lighting, and were supplied daily with sterile water only. Photographs were taken at regular intervals, and the occurrence of sporulation, sclerotium formation, or other morphogenetic changes was recorded for each plasmodium.

### (d) Total RNA Extraction and Sequencing

Total RNA extraction and RNA sample preservation were conducted in accordance with a previously published protocol [19,20]. Plasmodial tissue (5 mm^2^) was scraped from nutrient-free agar using a sterile toothpick while avoiding residual oats, then immersed in lysis buffer (6 M guanidinium in 1% Triton X-100). Samples were vortexed for 30 s at room temperature, rested for 1 min, and centrifuged at 16,000 × g for 1 min to remove debris. The supernatant was transferred to a fresh tube, mixed with an equal volume of 100% ethanol, and purified with the Direct-zol RNA MicroPrep Kit (Zymo Research) according to the manufacturer’s instructions.

RNA purity was assessed with a NanoDrop spectrophotometer (acceptance criteria: OD260/280≥ 1.8, OD260/230 ≥ 1.0). Concentration was measured using a Qubit RNA BR Assay, and integrity was verified on an Agilent TapeStation; only samples with RIN ≥ 8.0 were processed further.

For de novo transcriptome assembly, poly-A^+^ RNA was isolated with the NEBNext Poly(A) mRNA Magnetic Isolation Module and libraries were prepared using the NEBNext Ultra II Directional RNA Library Prep Kit for Illumina. Paired-end 150 bp sequencing (6 Gb, ∼40 million reads per sample) was performed on an Illumina NovaSeq X Plus.

For differential gene expression analysis, total RNA was processed with the same poly-A selection module and the NEBNext Ultra II RNA Library Prep Kit for Illumina, followed by single-end 75 bp sequencing on an Illumina NextSeq 500 (75-cycle High Output kit).

### (e) De novo Transcriptome Assembly

RNA-seq libraries derived from two experimental conditions—dark/nutrient-rich (DN) and light/starving (LS), each cultured for three days—were pooled for de novo transcriptome assembly. Raw reads were inspected with FastQC v0.12.1 for quality and adaptor contamination; problematic datasets were manually curated as needed. Assembly was performed with Trinity v2.1.1 [21], combining DN and LS reads.

Assembly quality was assessed with seqkit v2.6.1 (N50 statistics) and BUSCO v5.2.2 (dataset eukaryota_odb10) for completeness. Potential contamination was evaluated using BlobTools v1.1.1: contigs were taxonomically assigned by DIAMOND BLASTX v2.1.6 against UniProt reference proteomes, and GC-content plots were examined. Contigs forming distinct GC-content clusters and showing prokaryotic BLAST hits were flagged as putative contaminants, removed manually, and the assembly metrics were recalculated.

Functional annotation was generated by retrieving UniProt accessions from BLASTX-positive contigs and mapping them to protein descriptions and GO terms via the UniProt ID Mapper.

### (f) ifferential gene expression analysis

Transcript abundance was quantified as TPM using kallisto v0.51.1, with the de novo transcriptome assembly as the reference index. Differentially expressed genes (DEGs) were identified in edgeR v3.42.4 under a false-discovery-rate threshold of FDR < 0.05 and |log_2_FC| > 2. Hierarchical clustering of DEGs was performed with the pheatmap package, and all statistical analyses were executed in R (base v4.3.3).

## Result

### (a) Light- and Starvation-Induced Sporulation in *P. roseum* and *P. rigidum*

Previous studies showed that *P. polycephalum* achieves 100% sporulation after three days of starvation followed by light exposure (4/4 plasmodia in Renzel 2000 [7]; 40/40 in Chapman 1982 [18]). Applying the same regimen, we monitored six plasmodia each of *P. rigidum* and *P. roseum* for seven days. In *P. rigidum*, four of six samples sporulated—one on day 4, two on day 5, and one on day 6—while two failed to do so. In *P. roseum*, none of the six plasmodia formed fruiting bodies within seven days, and three specimens had died by day 7 (Figure1).

**Figure 1.**
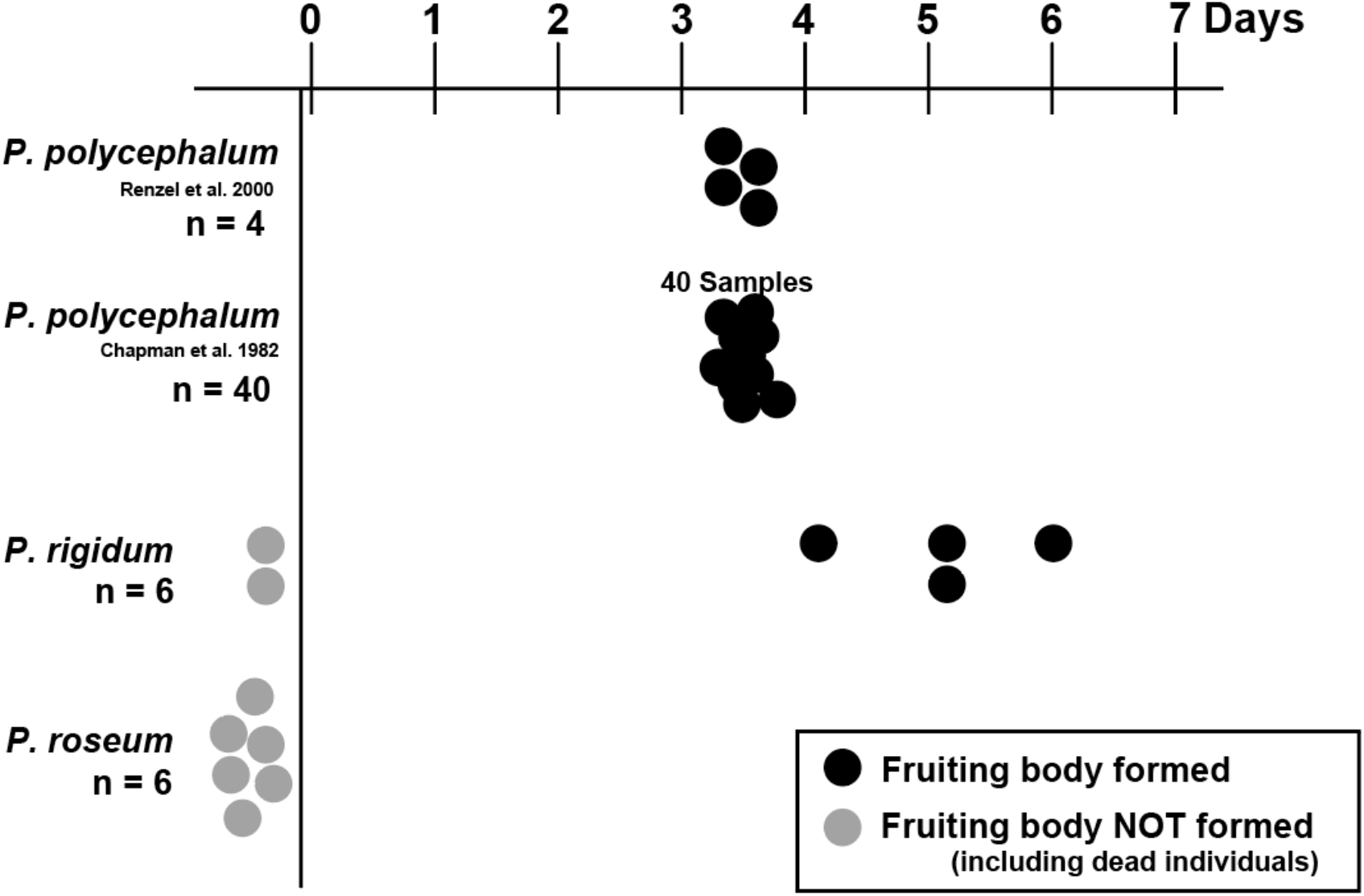
Time to Sporulation under Starvation + Light in Three *Physarum* Species. Black symbols indicate plasmodia that formed fruiting bodies; grey symbols indicate those that failed to sporulate within seven days. All *P. polycephalum* samples completed sporulation by day 3, whereas no *P. roseum* samples sporulated during the seven-day observation period. In *P. rigidum*, approximately 60% of plasmodia formed fruiting bodies by day 6.

### (b) De novo Transcriptome Assembly of *Physarum roseum*

Because *P. roseum*, unlike *P. polycephalum*, did not form fruiting bodies after starvation combined with light exposure, we next sought to characterise its transcriptomic response at this stage. Given the absence of a genome or transcriptome reference for this species, we first constructed a de novo transcriptome assembly.

Total RNA was extracted from plasmodia cultured under two contrasting conditions: dark/nutrient-rich (DN) and light/starved (LS). The DN condition represents routine maintenance, whereas LS is its physiological opposite. Paired-end 150 bp sequencing on an Illumina platform yielded 11.1 Gb of data. Assembly with Trinity produced 150,795 contigs and a BUSCO completeness of 94.1%. Because a previous study assembled RNA from several life cycle stages and obtained roughly 770 k contigs [21], the smaller contig number recovered in our work, based only on two plasmodial conditions, can be regarded as appropriate. Approximately 50% of contigs received UniProt-based functional annotations. Contamination screening by DIAMOND BLASTX against UniProt reference proteomes and visualisation with BlobTools revealed no GC-content clusters indicative of foreign sequences (Figure 2).

**Figure 2.**
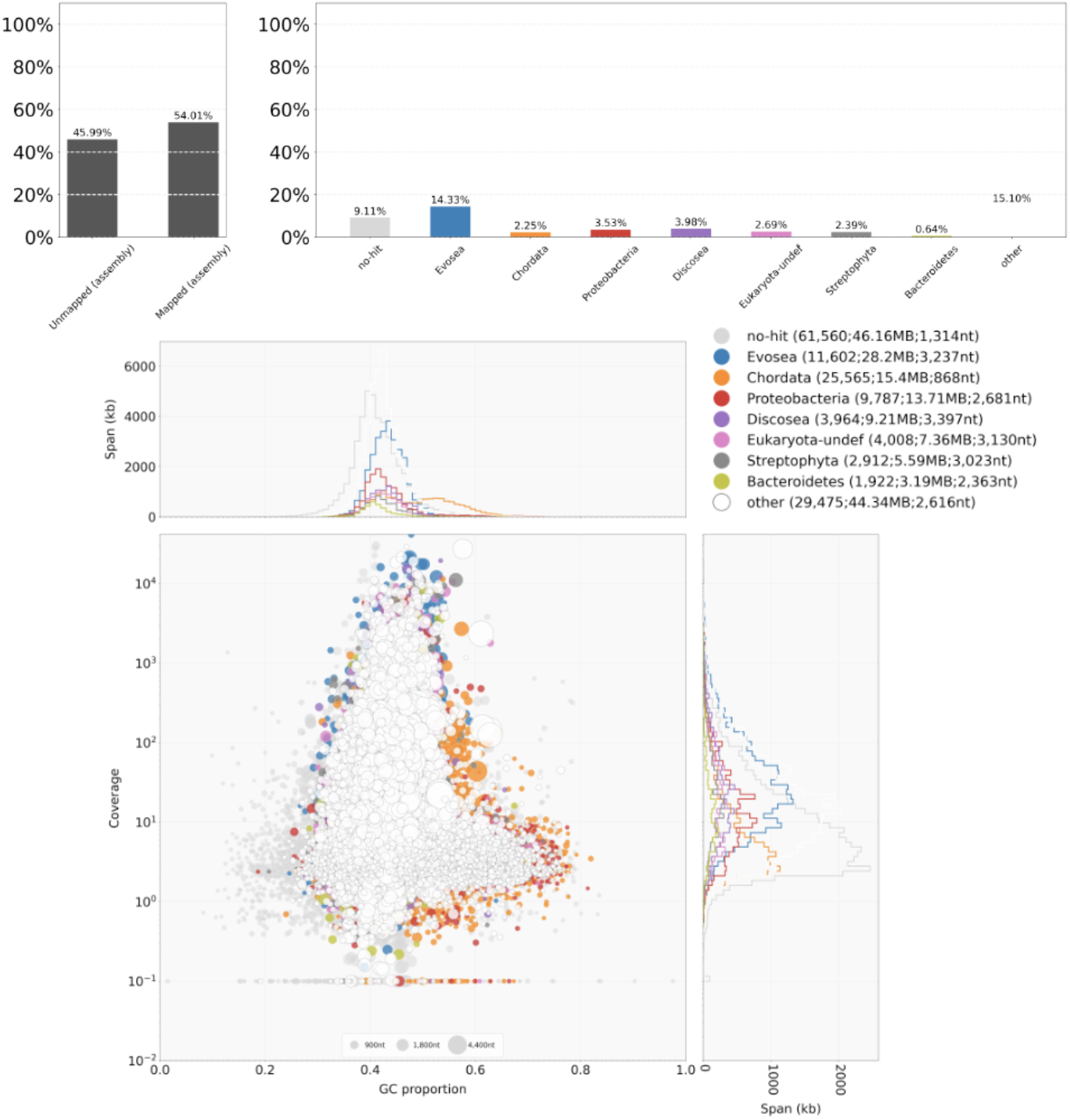
Contamination assessment with BlobTools: the analysis displays the relative abundance of each taxon present in the sequence reads together with their GC content. The “no-hit” reads match Evosea in GC profile, consistent with Myxomycetes, and thus likely originate from under represented Myxomycetes genomes rather than external contaminants.

### (c) Differential Gene Expression Analysis

To elucidate intracellular responses in *P. roseum*, we cultured plasmodia under four combinations of light (L, light; D, dark) and nutrition (S, starving; N, nutrient-rich): dark + starved (DS), light + starved (LS), dark + nutrient-rich (DN) and light + nutrient-rich (LN). Triplicate RNA-seq libraries (n = 3) were prepared for each condition, generating 10.5–30.7 million reads per sample that were mapped to the *P. roseum* reference transcriptome.

The mapping rates for all 12 libraries were all above 90%, with averages by condition of DN 93.1%, DS 98.6%, LN 95.2%, and LS 95.4%. These high matching rates suggest that de novo transcriptome assembly comprehensively captured expressed transcripts. Therefore, the analyse below will focus on the results of the gene expression variation analysis and their biological implications.

#### Global expression patterns

Spearman rank correlations showed that the DN condition—standard plasmodial maintenance— shared only modest similarity (ρ ≒ 0.7) with each of the other three conditions, whereas DS, LS, and LN were more closely related to one another (ρ = 0.8–0.9). Thus, both light exposure and nutrient depletion provide stimuli sufficient to shift the overall transcriptional profile away from that of standard.

#### DEG counts

We quantified genes exhibiting statistically significant expression changes as differentially expressed genes (DEGs) to obtain a numerical measure of their impact on the overall transcriptional profile. Under the DN condition, *P. roseum* exposed to nutritional starvation displayed more than 300 DEGs, and this response occurred irrespective of light stimulation (Figure3*a*). Interestingly, the set of genes influenced by switching between light and dark was far smaller—ranging from only a few dozen to none at all—when compared with the effect of nutrient status (figure3*a*). In a previous study, *P. polycephalum* responded to light stimulation with 6,985 DEGs [23]; hence *P. roseum* can be considered strikingly insensitive to light in comparison. Although starvation elicited more DEGs than light in *P. roseum*, the magnitude of this response does not approach the dramatic, system-wide expression changes reported during sporulation in earlier work.

**Figure 3.**
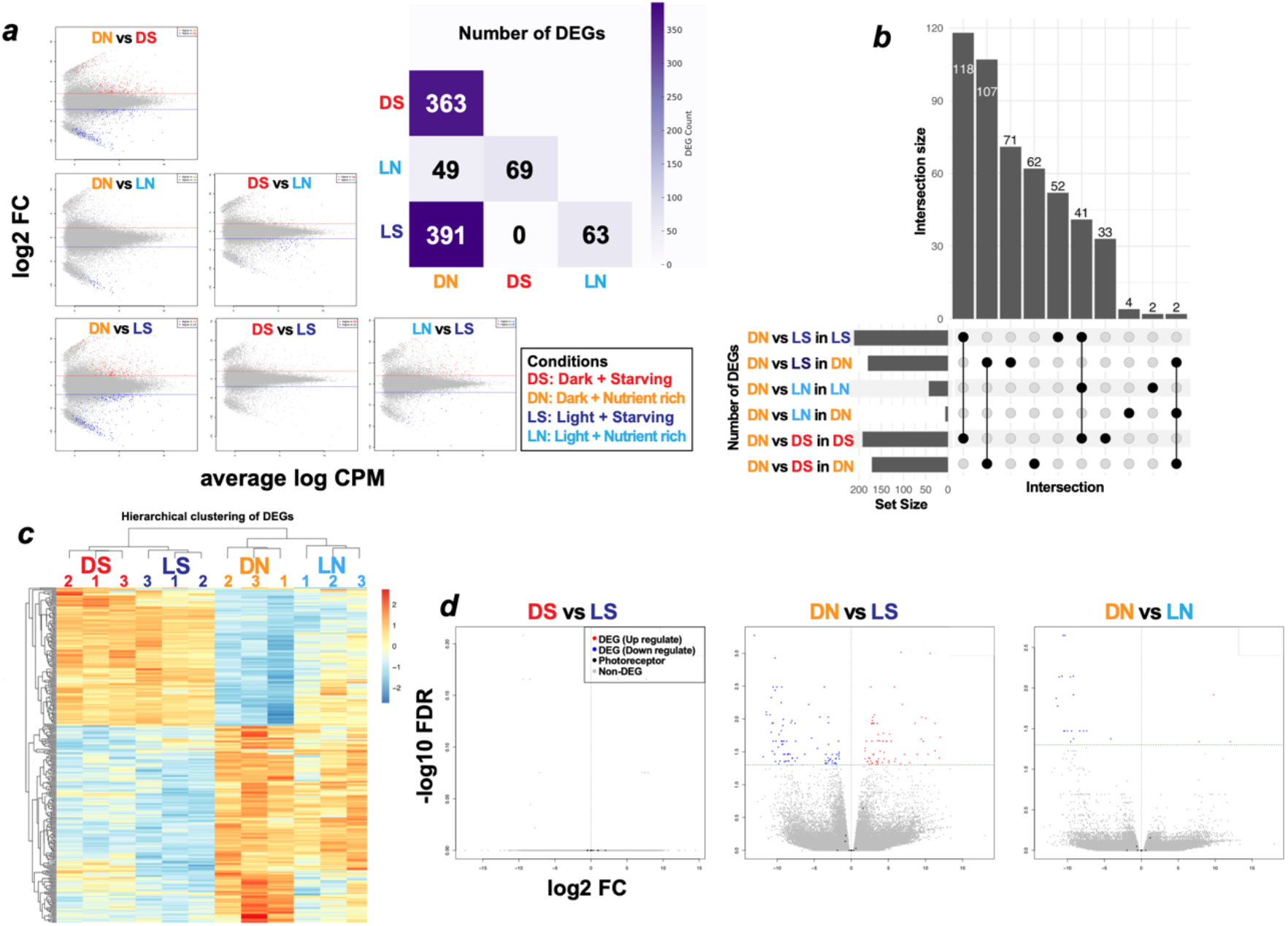
Results of differential gene-expression analysis. (*a*) DEG counts and MA plots across conditions. Relative to the dark, nutrient-rich baseline (DN), significant DEGs were detected in all other conditions, with the largest numbers arising from nutrient-deprived treatments. Light–dark comparisons (L vs D) yielded only limited DEG sets, indicating that most transcriptional changes are driven by nutrient availability (N vs S) rather than illumination. (*b*) UpSet plot of DEG overlap. Light–dark contrasts (L vs D) produce few DEGs and minimal overlap, whereas nutrient contrasts (N vs S) share extensive sets of consistently up- or down-regulated genes. A small subset is up-regulated in every comparison except the DN baseline, underscoring nutrient-dominated transcriptional programmed. (*c*) Hierarchical clustering of DEGs. Samples cluster cleanly by experimental condition, with nutrient availability emerging as the primary driver of grouping. (*d*) Volcano plot for light versus dark. Red points represent genes up-regulated under light condition, blue points those up-regulated under dark condition. Black points denote photoreceptor-related genes (isoforms collapsed); none meet the DEG threshold, and most exhibit minimal expression change.

#### Nutrient-driven signature

We next compared functional overlap among DEGs when the standard dark-nutrient (DN) culture condition was perturbed either by illumination or by nutrient withdrawal. Focusing on the two contrasts that yielded the largest DEG sets—DN vs LS and DN vs DS—more than 100 genes were commonly up-regulated by starvation regardless of light regime. Likewise, over 100 genes were down-regulated under those same conditions, meaning that roughly two-thirds of all DEGs were shared between the two comparisons (Figure3*b*). In other words, once *P. roseum* experiences any perturbation of the usual culture condition, the vast majority of expression changes are driven by nutrient status, while light responsiveness remains a minority component.

This pattern was mirrored in the expression-level clustering analysis. Hierarchical clustering of DEGs expressed in all samples (TPM > 0) revealed that the largest clade segregated strictly by nutrient availability (Figure3c). No clusters were intermixed across environmental conditions, further demonstrating that nutrient deprivation exerts a far stronger influence on the transcriptome of *P. roseum* than the light–dark regime.

#### Photoreceptor genes

Previous studies on *P. polycephalum* have identified two classes of photoreceptor-related genes: (i) a starvation-induced, phytochrome-like photoreceptor (*phyA, phyB*) [6, 21] and (ii) three genes that are constitutively expressed and were identified in the genome—the LOV domain photoreceptor *LovA*, the cryptochrome *cryA*, and a DNA photolyase (*plyA*) [21]. Accordingly, we investigated whether the light insensitive *P. roseum* exhibits comparable photoreceptor behaviour. First, none of the five photoreceptor genes reported in *P. polycephalum* was expressed under any condition tested. To identify function-based homologues, we predicted domains for the five candidate genes with InterProScan and queried those domains against the Pfam annotations assigned to our de novo transcriptome assembly. For each of the five genes, we detected a single transcript that possessed an identical photoreceptor related domain. However, none of these transcripts were classified as DEGs under any condition, and all displayed TPM values below 10. It suggests that they were scarcely expressed and therefore unlikely to be functionally active. Secondly, to cast a wider net for photoreceptor related genes in *P. roseum*, we conducted a GO term-based search. As a result, 92 photoreceptor-related genes (including isoforms) were retrieved; however, most were not expressed, and only 13 exhibited appreciable TPM values. Of these, six genes were highly expressed (≥ 10th percentile) across all conditions. Nevertheless, every one of the 13 genes displayed only minor expression shifts between conditions (|log_2_FC| < 2), and no significant DEGs was detected (Figure3d). These results indicate that although *P. roseum* does react to starvation, it neither synthesizes photoreceptors as *P. polycephalum* does nor undergoes the light-triggered sporulation and large-scale transcriptional changes characteristic of that species. Moreover, the lack of light-dependent DEGs despite the presence of constitutively expressed photoreceptors suggests that these receptors play at most a limited role in sporulation for *P. roseum*. Collectively, our data show that *P. roseum* is far less responsive to environmental cues—particularly light—than *P. polycephalum*, implying poor conservation of the sporulation regulatory network within the genus *Physarum*.

## Discussion

### (a) Evaluation of the de novo Transcriptome Assembly

In this study, we generated a de novo *P. roseum* transcriptome through a simple workflow: RNA from field-collected plasmodia was isolated by basic mechanical disruption and column purification. Despite this minimal preparation, the assembly achieved high BUSCO completeness, showing that fresh plasmodia alone can furnish a reliable molecular reference without axenic culture or elaborate decontamination. BLAST-based taxonomic screening revealed contigs assigned to Proteobacteria (Figure2). When these prokaryotic sequences were removed, the Complete BUSCO score (eukaryota_odb10) dropped substantially from 94.1% to 90.6%. Because prokaryotes rarely harbour eukaryotic core genes, two scenarios are plausible: (i) horizontal gene transfer has introduced bacterial genes into the slime-mould genome, or (ii) mis-assembly has erroneously merged bacterial and Myxomycetes transcripts. Given that the libraries were poly-A-selected, large-scale bacterial contamination is unlikely, making the former explanation more probable; yet definitive discrimination is difficult in the absence of a contamination-free reference for *Physarum*. Consequently, the putative bacterial contigs were retained for downstream analyses.

This approach provides a quick, convenient path to new molecular references for slime moulds. Because most Myxomycetes still lack reference sequences, our study stands as a proof-of-concept for generating practical transcriptomes with minimal effort.

### (b) Comparative Responses to Starvation + Light among Three *Physarum* Species

Previous studies have shown that *P. polycephalum* sporulates almost invariably after 3 days of starvation followed by light exposure [7,18]. Under the same regimen we observed no sporulation in *P. roseum*, whereas *P. rigidum* sporulated in four of six plasmodia (Figure1). These results demonstrate pronounced interspecific differences in physiological responses within the genus: *P. rigidum* behaves more like *P. polycephalum*, while *P. roseum* is markedly refractory. Although *P. rigidum* has not yet been included in molecular phylogenetic studies, its ecological similarity to *P. polycephalum*— especially in sporulation and culture behaviour—suggests a closer relationship than that of *P. roseum*. Focusing on the non-responsive species, we examined the transcriptomic dynamics of *P. roseum* under light (L) or dark (D) and nutrient-rich (N) or starving (S) conditions. Field-derived plasmodia yielded within-condition correlations of r = 0.8–0.9—slightly lower than for axenic cultures—yet sufficient for DEG detection. Relative to the dark, nutrient-rich baseline (DN), every perturbation elicited transcriptional change, but the magnitude was far smaller than the dramatic shifts that accompany sporulation in *P. polycephalum* (Figure3*a*). Light alone produced only limited DEGs, and starvation did not induce the phytochrome-like photoreceptors or downstream signalling cascade characteristic of *P. polycephalum* [23]. Although both species express a set of constitutive photoreceptor genes, their repertoires differ, and those of *P. roseum* are expressed at low levels and show no light-dependent regulation (Figure3*d*), arguing against a major role in sporulation.

The findings above demonstrate that *P. roseum* exhibits far fewer responses to environmental cues than *P. polycephalum* and is especially robust against light stimuli. Unlike *P. polycephalum, P. roseum* seems to cope with environmental changes, even harsh ones such as sudden illumination or prolonged starvation, by remaining in the plasmodial state as far as possible.

Although we were unable to induce sporulation in *P. roseum* under our laboratory regimen, but fruiting bodies are observed in nature. Two possibilities therefore arise. First, the starvation treatment used here may have been insufficient as a pre-conditioning step; in the field, complete starvation for three days or more is unlikely for a predatory plasmodium. Second, additional cues beyond starvation and light—such as pH, humidity, or, most plausibly, temperature—may be required. Moreover, natural environments show considerable variability in both nutrient availability and lighting conditions. Our laboratory conditions may not completely replicate the complexity of cues necessary to trigger sporulation in *P. roseum. P. roseum* is frequently reported from warm regions of Asia, Central America, and South America [17,24,25], and most isolates, including the one used in this study, were collected in summer. By contrast, *P. polycephalum* is typically cultured and induced to sporulate at 20–25 °C. *P. roseum* may therefore require relatively high temperatures in addition to the broadly conserved starvation-and-light stimuli. Such weak conservation of a supposedly fundamental sporulation response exceeds previous expectations and will be important for future work on the diversity and systematics of Myxomycetes.

Notably, *P. polycephalum* has become increasingly difficult to find in the field [26], whereas *P. roseum* appears to be widespread and stable—perhaps owing to the environmental robustness revealed here. Although *P. polycephalum* remains the canonical model species because its culture conditions are well defined and molecular resources are well developed, its limited field availability has begun to hamper studies that require multiple isolates [27]. From the standpoint of versatility and environmental tolerance, the continued reliance on *P. polycephalum* as the sole model may be questionable. With further refinement of culture methods and the acquisition of genomic references, *P. roseum* and other species could well emerge as more suitable model organisms for future molecular research.

## Data accessibility

Raw data used in this research are available from PRJNA1253572.

## Funding

This research funds from Masason Foundation, Hokuto Foundation for Bioscience, Chugai Foundation for Innovative Drug Discovery Science, Yamagata Prefecture, Tsuruoka City, and Keio University Academic Development Funds (Individual research).

## Acknowledgement

We thank the Saitama Midori-no-mori Nature Park for their cooperation in sampling the strains and Maaya Domon and Yuki Takai for providing technical support.

## Ethics

Ethical approval was not required for this work.

## Authors’ contributions

M.M.: conceptualization, data curation, formal analysis, funding acquisition, investigation, methodology, resources, validation, visualization, writing original draft, writing review and editing; P.Y.: supervision, validation, writing review and editing; N.K.: conceptualization, funding acquisition, methodology, project administration, resources, supervision, validation, writing original draft, writing review and editing.

## Conflict of interest declaration

The authors declare no competing or financial interests.

## Notes

### Competing Interest Statement

The authors have declared no competing interest.

### Summary of Updates

Author name (Phillip Yamamoto) has been revised

